# Environmental RNAi-based reverse genetics in the predatory mite *Neoseiulus californicus*: towards improved methods of biological control

**DOI:** 10.1101/2021.06.25.450003

**Authors:** Noureldin Abuelfadl Ghazy, Takeshi Suzuki

## Abstract

The predatory mite *Neoseiulus californicus* (McGregor) (Mesostigmata: Phytoseiidae) has been commercialized by manufacturers in the pest control industry and is used worldwide as a natural enemy of spider mites. However, because its genome has not been sequenced, reverse genetics techniques that could be used to analyze gene function have not been established. Here we partially sequenced the gene that encodes the vacuolar-type H^+^-ATPase (V-ATPase), an ATP-dependent proton pump, in *N. californicus* (*NcVATPase*) and then conducted a functional analysis using environmental RNA interference (eRNAi) by orally administering sequence-specific exogenous dsRNA (dsRNA-*NcVATPase*) to larvae and adult females. The larvae treated with dsRNA-*NcVATPase* took longer to develop and had lower survivorship, fecundity, and offspring viability at the adult stage than those treated with a control dsRNA. Adult females treated with dsRNA-*NcVATPase* showed significant reductions in survival, fecundity, and prey consumption, and their endogenous gene expression level of *NcVATPase* was reduced by approximately 65% compared with the control. Our findings suggest that the *NcVATPase* gene, silencing of which inhibits feeding and reproduction, is an excellent biomarker for investigating the eRNAi mechanism in *N. californicus*. The highly efficient experimental system of eRNAi established in this study paves the way for applied research using eRNAi to enhance the predatory ability of *N. californicus*.

**Key message:**

- Environmental RNAi-inducing double-stranded RNAs have the potential to improve biological control as well as biopesticide applications.
- We investigated the efficacy of eRNAi against the predatory mite *Neoseiulus californicus*, a major natural enemy of spider mites.
- Oral administration of dsRNA targeting *NcVATPase* decreased the gene expression level, developmental time, survival, fecundity, and prey consumption.
- *Neoseiulus californicus*, which was found to have the high eRNAi effects, can be used as a model for the study on eRNAi-mediated improvement of biological control.

## INTRODUCTION

The vacuolar-type H^+^-ATPases (V-ATPases) are a family of ATP-dependent proton pumps that play a role in luminal acidification of the intracellular compartments in eukaryotic cells (Nishi and Forgac 2002). Acidic intracellular compartments are required for several cellular processes including receptor-mediated endocytosis, intracellular trafficking, protein processing and degradation, and coupled transport of substrates across membranes (Harvey 1992; Forgac 2007; Wieczorek et al. 2009; Maxson and Grinstein 2014). Structurally, V-ATPases are composed of two domains: (1) the integral membrane-embedded V_0_ domain composed of a variable number of subunits (a, c, d, and e in insects; a, d, e, c, c′, and c″ in yeast; a, c, c″, d, e, and Ac45 in higher eukaryotes) that is responsible for proton translocation across the membrane via a rotary mechanism, and (2) the peripheral V_1_ domain with eight different subunits (A–H) that is responsible for ATP hydrolysis (Merzendorfer et al. 2000; Nishi and Forgac 2002; Toei et al. 2010; Maxson and Grinstein 2014; Cotter et al. 2015). V-ATPases that are expressed at the plasma membrane acidify the extracellular microenvironment and energize transcellular and paracellular transport (Nishi and Forgac 2002; Forgac 2007; Maxson and Grinstein 2014). In insects, V-ATPases energize K^+^ pumps to maintain a higher concentration of potassium, thus creating an electrical difference that allows nutrient uptake by midgut cells, fluid secretion by Malpighian tubules, and fluid absorption by insect ovarian follicle cells (Klein 1992; Harvey et al. 1998; Wieczorek et al. 2000). In addition, V-ATPases are reported to expel H^+^ from cells in the insect midgut to create a membrane potential that drives Na^+^ that is linked to an amino acid into the cell via nutrient amino acid transporters (Harvey et al. 2009; Fu et al. 2014, 2015).

Post-transcriptional gene silencing mediated by RNA interference (RNAi) is a conserved biological process in eukaryotes that is triggered by sequence-specific double-stranded RNA (dsRNA) (Fire et al. 1998; Hannon 2002). Accumulating experimental evidence in arthropods shows that RNAi can be initiated by introducing exogenous dsRNA against the gene of interest through direct injection or through ingestion, either of an artificial diet or drinking water containing the dsRNA itself or of bacteria or plant cells that express the dsRNA (Baum et al. 2007; Khila and Grbić 2007; Huvenne and Smagghe 2010; Yao et al. 2013; Suzuki et al. 2017a; Sijia et al. 2019; Ghazy et al. 2020; Bensoussan et al. 2020). Because the RNAi method of gene silencing is versatile, simple, and effective, it has been employed in a vast array of research, from functional genomic studies to the potential application of spraying dsRNA as a bio-pesticide against pest arthropods. In the context of arthropod pests and their natural enemies, RNAi studies that investigate the “off-target” effects of dsRNA in a natural enemy of a target pest and the effect of manipulation of reproductive or behavioral gene expression in a predator or parasite can be applied to enhance their functions as a predator or parasite in biological control programs (Pomerantz and Hoy 2015).

The predatory mite, *Neoseiulus californicus* (McGregor) (Mesostigmata: Phytoseiidae) is widely used as a biological control against tetranychid mites (Prostigmata: Tetranychidae). As a predatory omnivore, *N. californicus* also feeds on other phytophagous mites, small insects, and pollen (Castagnoli and Simoni 2003; McMurtry et al. 2013). The ability of *N. californicus* to feed on alternative prey when tetranychids are scarce has advanced its role as an important biological control agent that is commercially produced by several companies worldwide. To date, only a few studies have reported the use of RNAi-based reverse genetics in phytoseiid mites. These studies have shown that, in a fashion similar to nematodes and other arthropods, the phytoseiid mites *Phytoseiulus persimilis* Athias-Henriot and *Galendromus* (=*Metaseiulus*) *occidentalis* (Nesbitt) are amenable to environmental RNAi (eRNAi) when it is induced by delivering exogenous dsRNA orally (Ozawa et al. 2012; Wu and Hoy 2014, 2016; Pomerantz and Hoy 2015; Pomerantz et al. 2015; Sijia et al. 2019).

The study described here is the first RNAi study in the genus *Neoseiulus*. We isolated and partially sequenced the gene encoding V-ATPase in *N. californicus* (*NcVATPase*; GenBank accession number: MK281632) and then analyzed the gene function by using an eRNAi method based on the oral delivery technique (Ghazy and Suzuki 2019). We found that the *NcVATPase* gene is involved in development during the immature stages, and in adult survival, reproduction, feeding, and appetite. This study focused on establishing an efficient method of orally feeding dsRNA to suppress the expression of a target gene linked to behavioral changes and thus paves the way for using eRNAi-based genetic screens to enhance the functions of predatory mites.

## MATERIALS AND METHODS

### Source of N. californicus colony

A commercial population of *N. californicus* was obtained from Sumika Technoservice (Takarazuka, Japan) in 2017 and has been routinely reared on detached kidney bean leaves (*Phaseolus vulgaris* L.) infested with the two-spotted spider mite (*Tetranychus urticae* Koch) as its prey. Mites were maintained and all experiments in this study were conducted at 25 °C, 50%–65% relative humidity and a light:dark cycle of 16:8 h. An air pump–based system was used to collect mites for experiments (Suzuki et al. 2017b).

### NcVATPase gene sequencing

Total RNA was extracted from ca. 400 adult females of *N. californicus* frozen in liquid nitrogen with NucleoSpin RNA Plus (Macherey-Nagel, Düren, Germany), following the manufacturer’s instructions. The quality and quantity of RNA were measured using a spectrophotometer (NanoPhotometer N60; Implen, Munich, Germany). cDNA was synthesized from 3 µg of total RNA using reverse transcriptase (SuperScript II Reverse Transcriptase; Thermo Fisher Scientific, Waltham, MA) and an oligo (dT)_12–18_primer (Thermo Fisher Scientific), according to the manufacturer’s protocol. cDNA was then stored at –30 °C until used. To amplify *NcVATPase*, degenerate primers (5′-GGCCACCATCCAGGTGtaygargarac-3′ and 3′-ccnccnctraaGAGGCTGGGGCACTG-5′) were designed based on aligned amino acid sequences of V-ATPase from the predatory phytoseiid mite *G. occidentalis* (Mesostigmata: Phytoseiidae) (GenBank accession number: XP_003741079.1) and the parasitic mite *Varroa destructor* Anderson and Trueman (Mesostigmata: Varroidae) (GenBank accession number: XP_022670783.1) by using the CODEHOP program (Rose et al. 1998). The resulting sequence of *NcVATPase* shares 85% similarity with that of *G. occidentalis* and 75% similarity with that of the *Varroa* species. An evolutionary analysis of *NcVATPase* was performed using MEGA 10.05 software on ClustalW-aligned amino acid sequences of V-ATPases from several mite, tick, and insect species by the neighbor-joining method and with 500 bootstrap replicates. The species list and GenBank accession numbers are presented in Table S1.

### dsRNA synthesis

The partial sequence of *NcVATPase* was used to design sequence-specific primers containing T7 promoter sequences (5′-TAATACGACTCACTATAGGGGGAAACCTCTCTCCGTCGAA-3′ and 5′-TAATACGACTCACTATAGGGGCCATCGAACTCTGTTTCCA-3′) to amplify a 344-bp DNA fragment of *NcVATPase*. For the negative control, a 659-bp DNA fragment of *DsRed2* was amplified from 1 ng of pDsRed2-N1 plasmid (Clontech, Mountain View, CA, USA) with primers (5′-TAATACGACTCACTATAGGGCGTGCACTCGTACACTGAGG-3′ and 5′-TAATACGACTCACTATAGGGTCATCACCGAGTTCATGCG-3′), according to the method used by Tokuoka et al. (2017). The PCR amplifications were carried out using a DNA polymerase (Phusion High-Fidelity DNA Polymerase; New England Biolabs, Hitchin, UK). The DNA fragments were then purified with a NucleoSpin Gel and PCR Clean-Up Kit (Macherey-Nagel). The integrity of the purified DNA fragments was confirmed by 2% (w/v) agarose gel electrophoresis, quantified with the spectrophotometer, and stored at -30 °C until used. Both types of DNA fragment (0.1 µg) were used as templates to synthesize RNA using an *in vitro* transcription kit (*in vitro* Transcription T7 Kit; Takara Bio, Shiga, Japan) in 1.5 mL centrifuge tubes. The fragments were denatured at 95 °C for 5 min then slowly cooled to room temperature to facilitate dsRNA formation (Suzuki et al. 2017a). The dsRNA was purified with a phenol:chloroform:isoamyl alcohol (25:24:1) solution and precipitated with ethanol. The dsRNA precipitate was dissolved in nuclease-free water, and then quantity and quality were checked on a spectrophotometer and by electrophoresis in 2% (w/v) agarose gel.

### RNAi bioassay

The *in vitro* droplet feeding method (Ghazy and Suzuki 2019) was used to deliver dsRNA-*NcVATPase* or dsRNA-*DsRed2* (negative control) to age-synchronized protonymphs and adult females of *N. californicus*. Newly hatched larvae (∼10-h old) or 2–3-day-old adult females were confined in 0.5-mL polypropylene tubes and a single 1-µL droplet of either type of dsRNA (1 µg µL^-1^) was placed on the inner surface of each tube lid. The test mites were allowed to feed on the droplet for 24 h at 25 °C. A Brilliant Blue FCF tracer dye was first added to the dsRNA solution so that successful delivery could be confirmed by the presence of the color blue in the alimentary canal of treated adult females and protonymphs that emerged from larvae during the 24 h after treatment (Fig. 1). As Ghazy and Suzuki (2019) note, the blue color of the tracer dye is not observable in the alimentary canal of larvae but, because at 25 °C the larval stage is usually shorter than 24 h (e.g., Gotoh et al. 2004), the color is visible in ∼100% of the protonymphs that emerge. After dsRNA delivery had been confirmed, the adult females were transferred to normal rearing conditions and provided with *T. urticae* at random life stages as prey on discs of kidney bean leaves (2.5 cm diameter, 2 to 5 females/disc). The adult females were observed for 10 days after treatment and their survival and fecundity were recorded every day. The feeding activity of adult females was also examined for three successive days after the dsRNA treatment. To do this, a known number of *T. urticae* eggs was provided to groups of two females of *N. californicus* every day on leaf discs (2 cm diameter) and the number of eggs consumed was recorded. As for protonymphs, after dsRNA delivery they were transferred individually onto leaf discs (1.5 cm diameter) with a surplus number of *T. urticae* at random life stages, and the developmental time to adulthood, survival at 10 days after treatment, fecundity (during the first 5 days of oviposition) of emerged adult females, and hatching success of their eggs were determined.

**Fig. 1.**
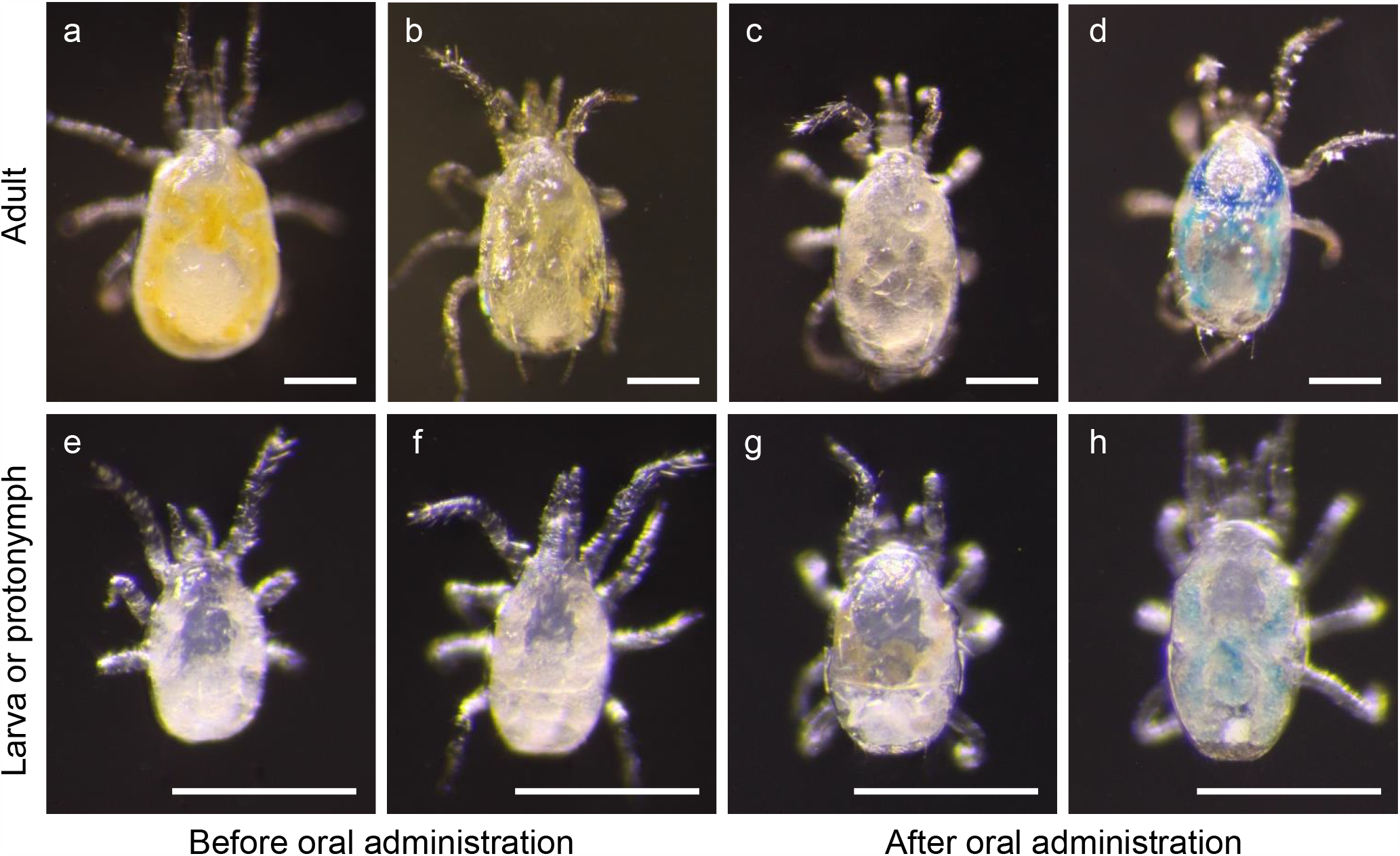
Images of *N. californicus* adult mites (**a**–**d**) and early immature mites (**e**–**h**) that illustrate the droplet feeding method. (**a**) An adult female mite just after collection and (**b**) after being starved for 24 h. Water was then provided, and images taken 24 h later show mites that ingested (**c**) water or (**d**) water containing a blue tracer dye (Brilliant Blue FCF). (**e**) A newly hatched larva (∼6 h old) that was confined for 24 h to ensure completion of the larval stage, and (**f**) a newly emerged protonymph, which will be able to ingest a test solution. (**g**) shows a protonymph that received water for 24 h from the larval stage and (**h**) one that received water containing a blue tracer dye. Scale bar: 100 µm.

### Real-time quantitative reverse transcription-PCR analysis

Adult females treated with dsRNA-*NcVATPase* or dsRNA-*DsRed2* were collected at 4 days after treatment (∼50 females in each of three biological replicates). Total RNA was extracted using NucleoSpin RNA Plus (Macherey-Nagel), and single-stranded cDNA was synthesized by reverse transcription of total RNA using a High Capacity cDNA Reverse Transcription Kit (Applied Biosystems, Foster City, CA, USA). Real-time quantitative reverse transcription PCR (real-time RT-qPCR) reactions were performed in 3 technical replicates for each sample with Power SYBR Green Master Mix (Applied Biosystems). The real-time RT-qPCR reactions were performed on an ABI StepOnePlus Real-Time PCR System (Applied Biosystems). The *NcVATPase* primers used for real-time RT-qPCR were 5′-GGAGTTCAATCCCTCCAACA-3′ and 5′-GGGCTCAAGCATGATTTTGT-3′. A fragment of *N. californicus Actin* gene was amplified using primers (5′-TGAGGCAATCGGTGTGTTTG-3′ and 5′-TTTTCACGATTGGCCTTGGG-3′) designed from the nucleotide sequence of the *Actin 1* gene in the predatory mite *N. cucumeris* (GenBank accession number: KC335208.1). The nucleotide sequence of the fragments amplified from *N. californicus* (GenBank accession number: MK848403) has 95% similarity with that of *Actin 1* in *N. cucumeris*. The real-time RT-qPCR was performed using primers specific to *N. californicus Actin* (5′-TGGCACCACACCTTCTACAA-3′ and 5′-GGGTTTTCACGATTGGCCTT-3′). The expression level of the *Actin* gene was used as a reference gene when normalizing the expression data of *NcVATPase*. Amplification efficiencies for the target gene (E_T_) and reference gene (E_R_) were 98.1% and 106.1%, respectively. The average threshold cycle (Ct) value was calculated from three technical replicates of each biological replicate. The expression value of the target gene (T) was normalized to the reference gene (R), and the normalized relative quantity (NRQ) was calculated as follows: NRQ = (1 + E_R_)^CtR^ / (1 + E_T_)^CtT^.

### Data analysis

The survival curves were plotted using the Kaplan–Meier method (R function: survfit, package: survival), and survival curves were compared using the log-rank test (R function: survdiff, package: survival). Differences in fecundity, prey egg consumption, and the relative quantity of *NcVATPase* gene expression between the dsRNA treatments in mites that had been treated when they were adult females, and in developmental time to adulthood and fecundity in mites that had been treated while they were protonymphs were statistically analyzed with a *t*-test (R function: t.test). A chi-square test was performed to analyze the survival of mites treated when they were protonymphs and the hatchability of eggs from females that emerged. Data were analyzed and visualized with R v. 3.5.1 (R Core Team, 2020).

## RESULTS

### Sequence analysis of *NcVATPase*

The partial sequence fragment of *NcVATPase* (419 bp) encoded 123 amino acids and had a molecular weight of 13.86 kDa. The amino acid sequence has high similarity with V-ATPase A from *G. occidentalis* (93%), *V. destructor* (83%; GenBank accession numbers: XP_022670784.1 and XP_022670783.1), the ectoparasitic mite *Tropilaelaps mercedesae* (Mesostigmata: Laelapidae) (80%; GenBank accession number: OQR76956.1), and the deer tick *Ixodes scapularis* (Ixodida: Ixodidae) (72%; GenBank accession number: XP_029849202.1). The phylogenetic tree constructed by using the neighbor-joining method from V-ATPases in mites, ticks, and several species of insects (Table S1) showed that *N. californicus* V-ATPase was clustered with those of mesostigmatid mites: *G. occidentalis, T. mercedesae*, and *V. destructor* (Fig. 2).

**Fig. 2.**
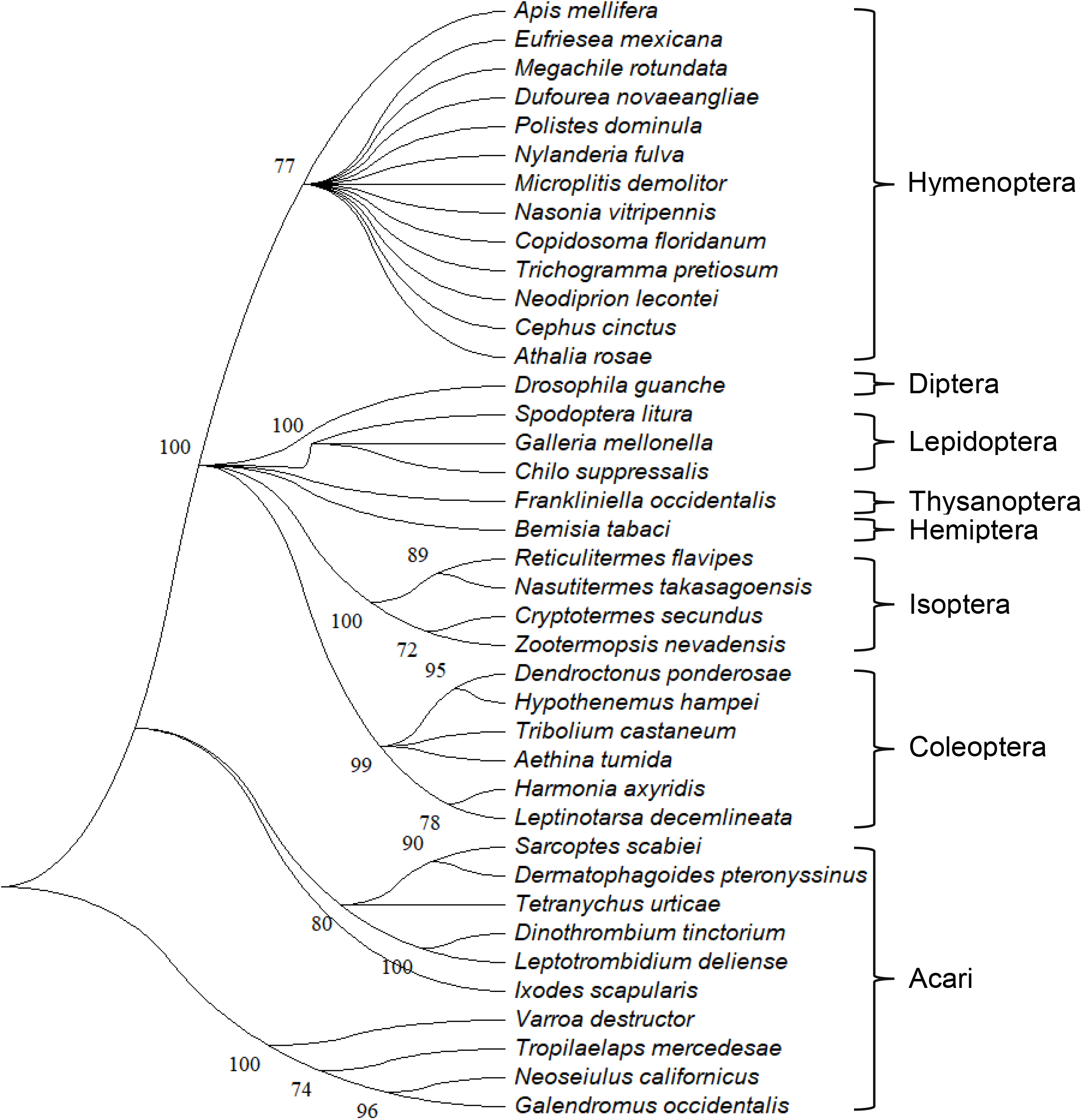
A phylogenetic tree of the *V-ATPase subunit A* gene. The tree was constructed by the neighbor-joining method with 500 bootstrap replicates generated from protein sequence alignments of *V-ATPase subunit A* in *N. californicus* with those in several mite, tick, and insect species. GenBank accession numbers for the species used in the analysis are listed in Table S1. The numbers on interior branches are bootstrap values.

### RNAi in adult females

The expression level of endogenous *NcVATPase* transcripts was significantly lower in adult females of *N. californicus* treated with dsRNA-*NcVATPase* and was 35% of that in the control (Fig. 3a). Oral delivery of dsRNA-*NcVATPase* significantly reduced mite survival to 25% at 10 days after treatment (Fig. 3b). Mites treated with dsRNA-*NcVATPase* ceased producing eggs within 3 days after treatment and did not recover reproductive ability even at 10 days after treatment. In total, mites treated with dsRNA-*NcVATPase* produced up to 15 times fewer eggs than *DsRed2*-treated control mites (Fig. 3c). Furthermore, mites treated with dsRNA-*NcVATPase* were less voracious, with 22% food consumption (8.7 prey eggs/female/3 days) relative to the control (40.4 prey eggs/female/3 days) (Fig. 3d). In addition to reductions in level of *NcVATPase* transcript, survival, fecundity, and feeding, a small body size was observed as a typical phenotype in mites treated with dsRNA-*NcVATPase* (Fig. 3e).

**Fig. 3.**
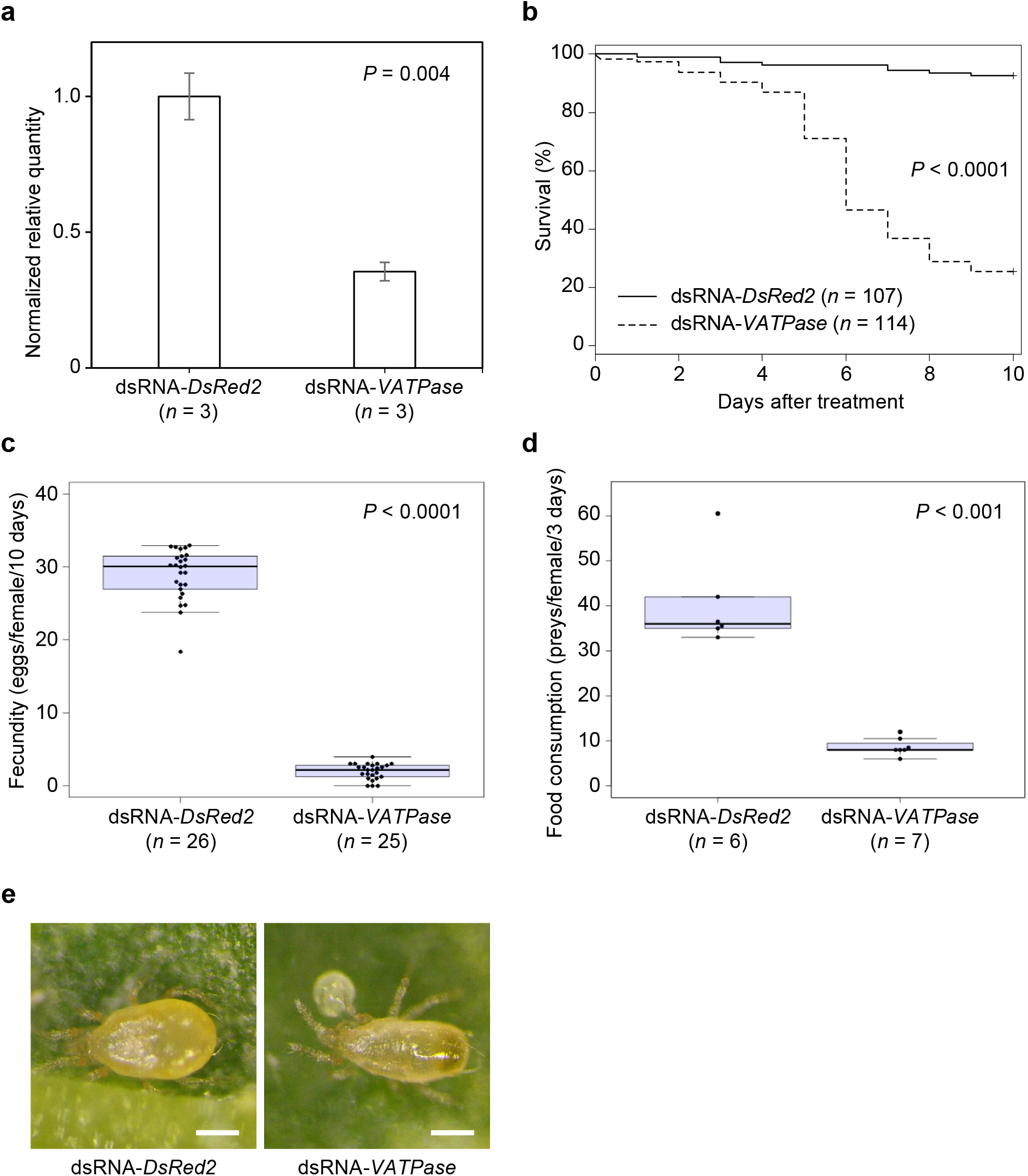
The effect of 1-µg µL^-1^ dsRNA-*NcVATPase* or dsRNA-*DsRed2* (negative control) delivered by the droplet feeding method on the endogenous *NcVATPase* gene expression levels, survivorship, fecundity, and food consumption in adult *N. californicus* females at 25 °C. (**a**) *NcVATPase* gene expression in adult females relative to the expression of the actin reference gene at 4 days after treatment with dsRNA-*NcVATPase* and dsRNA-*DsRed2*. Data are represented as the mean ± SE and compared using the *t*-test. (**b**) Survivorship of adult females for 10 days after treatment with dsRNA-*NcVATPase* and dsRNA-*DsRed2*. Survival curves were plotted by the Kaplan–Meier method and compared using the log-rank test. (**c**) Fecundity of adult females for 10 days after treatment with dsRNA-*NcVATPase* and dsRNA-*DsRed2*. (**d**) The average number of prey eggs (*T. urticae*) consumed per female *N. californicus* in three successive days. (**e**) Images show the body phenotype associated with each dsRNA treatment. Scale bar: 100 µm. Smaller bodies were observed in 100% of females given dsRNA-*NcVATPase* treatment. (**a, b, c, d**) Data are from at least three independent experimental runs. (**c, d**) Data are represented by bee-swarm and box-and-whisker plots and compared using a *t*-test. The central line (second quartile, Q2) indicates the median, the distance between the bottom (first quartile, Q1) and top (third quartile, Q3) of the box indicates the interquartile range (IQR), and the bottom whisker and top whisker indicates the minimum and maximum value, respectively. Outliers that are outside the range between the lower [Q1 - 1.5 × IQR] and upper limits [Q3 + 1.5 × IQR] are plotted outside of the IQR.

### RNAi in larvae

Within 24 h of being confined, the larvae became protonymphs in which the alimentary canal was colored with the blue tracer dye, thus providing evidence that dsRNA delivery had been successful. The time needed for larvae treated with *NcVATPase* to reach the adult stage was only slightly longer than that for the control, but the difference was nevertheless significant (Fig. 4a). The survival rate of mites that emerged from larvae treated with dsRNA-*NcVATPase* was 78% (*n* = 54), which was significantly lower than in control mites (92%, *n* = 56, Fig. 4b). The fecundity of adult females that emerged from larvae exposed to dsRNA-*NcVATPase* was significantly lower than in the control (Fig. 4c), and the hatchability of their eggs was also significantly lower, with many of the eggs deformed (Figs. 4d, e).

**Fig. 4.**
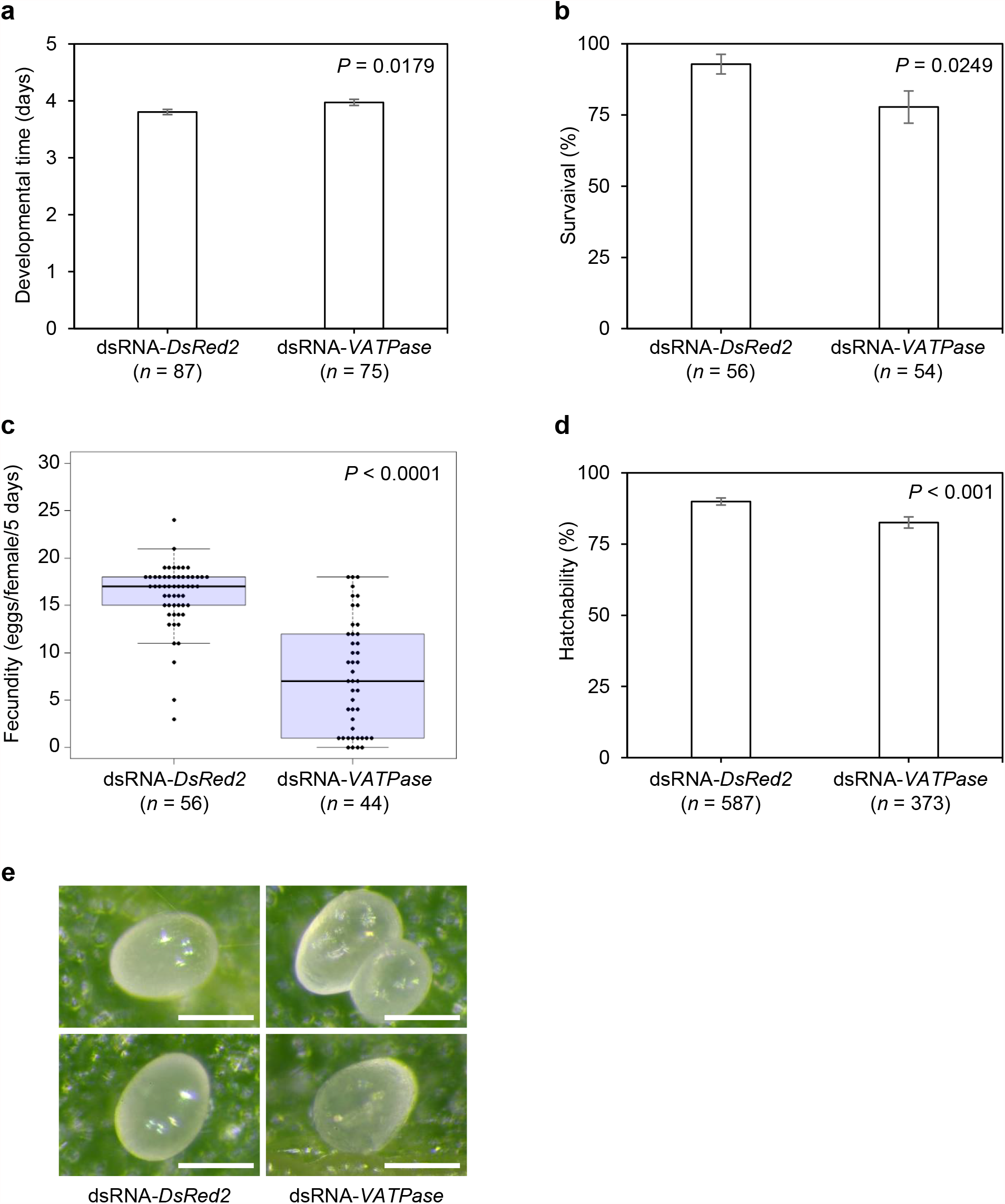
The effect of 1 µg µL^-1^ dsRNA-*NcVATPase* or dsRNA-*DsRed2* (negative control) delivered to *N. californicus* larvae by the droplet feeding method on the development, survival, and fecundity in adult females at 25 °C. (**a**) Time taken from larval stage to adulthood after treatment with dsRNA-*NcVATPase* and dsRNA-*DsRed2*. (**b**) Survival at 10 days after treatment of females that emerged from treated larvae. (**c**) Fecundity of adult females that developed from larvae treated with dsRNA-*NcVATPase* and dsRNA-*DsRed2*, measured during the first 5 days of oviposition. Data are represented by bee-swarm and box-and-whisker plots and compared using a *t*-test. (**d**) Hatchability of eggs produced by adult females of *N. californicus* that had been exposed when larvae to dsRNA-*NcVATPase* and dsRNA-*DsRed2*. (**a, b, d**) Data are represented as percentage ± SE and compared using the chi-square test. Numerical values in parentheses indicate the total number of mites tested. (**e**) Images show the deformed shape and soft shell of the egg phenotype associated with dsRNA-*NcVATPase* treatment. Scale bar: 100 µm. **(a, b, c, d**) Data are from at least three independent experimental runs.

## DISCUSSION

In this study, we partially sequenced the gene that encodes V-ATPase subunit A in the predatory mite *N. californicus*. The amino acid sequence from a retrieved DNA fragment of *NcVATPase* showed 72% to 83% similarity with *V-ATPase* genes from *G. occidentalis, V. destructor*, the ectoparasitic mite *T. mercedesae*, and the deer tick *I. scapularis*. A phylogenetic analysis of the V-ATPase amino acid sequences in known mites, ticks, and several insect species revealed that the V-ATPase of *N. californicus* belongs to a cluster found in Mesostigmata mites (Fig. 2). A functional analysis of *NcVATPase* was performed by using an eRNAi technique in which dsRNA was orally delivered (Ghazy and Suzuki 2019). We found that the *NcVATPase* gene is involved in multiple functions, and its silencing affected development, survival, fecundity, and egg maturation in *N. californicus*. The results are generally consistent with previous reports on insects, ticks, mites, and nematodes and indicate that the functions of *V-ATPase* in invertebrates are conserved (Huvenne and Smagghe 2010; Knight and Behm 2012; Petchampai et al. 2014; Suzuki et al. 2017a; Bensoussan et al. 2020; Ghazy et al. 2020).

Oral delivery of dsRNA-*NcVATPase* to adult *N. californicus* females resulted in significant reductions in mRNA transcript level, mite survival, fecundity, and feeding appetite (Fig. 3). These results are compatible with those of a previous study on *T. urticae*, which reported that targeting the *TuVATPase* gene with dsRNA significantly reduces adult survival and fecundity (Suzuki et al. 2017a). The reduction in mite survival and fecundity when the *V-ATPase* gene was silenced (Figs. 3b, c) may be correlated with the lower predation activity of *N. californicus*, measured as the number of prey eggs consumed (Fig. 3d). Although the mechanism involved is still unknown, it has been reported that gene silencing of *V-ATPase* decreases insects’ appetite. For instance, co-silencing the *coatomer β* and *V-ATPase A* genes reduces the feeding activity of the cotton bollworm *Helicoverpa armigera* (Hübner) (Lepidoptera: Noctuidae) (Mao et al. 2015). After dsRNA-mediated silencing of *V-ATPase* genes, feeding is also inhibited in the tomato leafminer *Tuta absoluta* (Meyrick) (Lepidoptera: Gelechiidae) and the western corn rootworm *Diabrotica virgifera virgifera* LeConte (Coleoptera: Chrysomelidae) (Baum et al. 2007; Camargo et al. 2016.). In our study, the inhibited feeding may have caused nutritional deficiency in adults, which in turn hindered egg production and/or embryonic development and resulted in lower fecundity being observed after the treatment with dsRNA-*NcVATPase* (Fig. 3b).

When mites were treated from the larval stage, they took significantly longer to develop (Fig. 4a) and survived for a shorter time (Fig. 4b). In addition, even after they emerged as adults, the treated mites still had phenotypes related to *V-ATPase* gene silencing, such as reduced fecundity and production of deformed immature eggs with low hatchability (Figs. 4c– e). Transgenerational RNAi is often called “parental RNAi” and has been reported in nematodes, insects, and mites (e.g., Fire et al. 1998; Bucher et al. 2002; Khila and Grbic 2007; Wu and Hoy 2014); however, in most of these studies the dsRNA was injected into or ingested by adults or pupae. In our study, we showed that the effects of orally administering dsRNA in the early immature stages of *N. californicus* persisted into the adult stage and even into the next generation, which indicates the robust and persistent effect of parental RNAi in this species. Hoy et al. (2016) have suggested that systemic and parental RNAi by oral administration of dsRNA in *G. occidentalis* is mediated by the *clathrin heavy chain* gene, which is responsible for endocytosis of dsRNA uptake, and by the *RNA-dependent RNA polymerase* gene (of which at least three copies are present in this species), which is responsible for dsRNA amplification. The robust and long-term parental RNAi effects in *N. californicus*, a member of the same Mesostigmata suborder, also suggest the existence of a similar RNAi machinery and pathway to that seen in *G. occidentalis*.

The downregulation of the *V-ATPase* genes in insects and mites has been frequently reported to cause growth inhibition, molting defects, and reproductive disorders. For instance, nymphal molting defects and growth inhibition result from RNAi-mediated gene silencing of *V-ATPase subunit B* in the smokybrown cockroach *Periplaneta fuliginosa* (Serville) (Blattodea: Blattidae) or of *V-ATPase subunit H* in the migratory locust *Locusta migratoria* L. (Orthoptera: Acrididae) (Li and Xia 2012; Sato et al. 2019). Similar phenotypes have also been observed in the small hive beetle *Aethina tumida* Murray (Coleoptera: Nitidulidae) when the *V-ATPase subunit A* gene was silenced (Powell et al. 2017). Decreased fecundity is the most prominent phenotype induced by RNAi-mediated silencing of the *V-ATPase* gene in *T. urticae* (Suzuki et al. 2017a; Bensoussan et al. 2020; Ghazy et al. 2020). In our study, the adults that emerged from treated larvae produced deformed and immature eggs, which may indicate that nutrient uptake or nutrition utilization was interrupted in the treated mites. Yao et al. (2013) reported that gene silencing of *V-ATPase subunit B* or *subunit D* in the corn planthopper *Peregrinus maidis* (Ashmead) (Hemiptera: Delphacidae) has a negative impact on nymphal development, survival, and fecundity and inhibits the development of the reproductive organs in treated females. They further reported that nymphs injected with dsRNA of *V-ATPase subunit B* or *subunit D* lose their ability to produce eggs as adult females. This may indicate that V-ATPases play a role in yolk processing during embryogenesis, as observed in the fruit fly *Drosophila melanogaster* Meigen (Bohrmann and Braun 1999). Against the background of this previous evidence, the results of our study further highlight that the *V-ATPase* genes have diverse and conserved functions in eukaryotes.

To the best of our knowledge, this is the first report on the response to RNAi in *N. californicus*. Because the oral method used to deliver dsRNA is simple and effective, it can be used for a vast array of RNAi-based gene function studies. The amenability to RNAi displayed by *N. californicus* in this study and by other phytoseiid mites, such as *P. persimilis* (Ozawa et al. 2012; Sijia et al. 2019) and *G. occidentalis* (Wu and Hoy 2014, 2016; Pomerantz and Hoy 2015; Pomerantz et al. 2015), will pave the way for more in-depth investigations into phytoseiid genetics and will facilitate the study of RNAi off-target effects, if any, in this highly important group of natural enemies. In addition, the robust and long-term RNAi effects observed in *N. californicus* indicate that dsRNA shows potential for use as an oral supplement to enhance predation performance. Based on the reverse genetic methodology developed in this study, we will conduct a highly efficient eRNAi screen to identify target genes that regulate feeding behavior in *N. californicus*.

## ACKNOWLEDGEMENTS

This study was supported by the following grants to TS: JSPS KAKENHI (18H02203 and 21H02193), JST OPERA (JPMJOP1833), and from the Bio-oriented Technology Research Advancement Institution (JPJ009237), as part of the Moonshot Research and Development Program, “Technologies for Smart Bio-industry and Agriculture” run by the Ministry of Agriculture, Forestry and Fisheries. NAG was supported by JSPS Invitational Fellowships for Research in Japan (L19542).

## AUTHOR CONTRIBUTIONS

NAG and TS conceived and planned the study. NAG performed experimental procedures and collected data. NAG and TS performed analysis and wrote the manuscript.

## CONFLICTS OF INTEREST

The authors declare no conflicts of interest.

## FIGURE LEGENDS

**Table S1** Species list and GenBank accession numbers of species used in the evolutionary analysis of *N. californicus V-ATPase*.

## Notes

### Competing Interest Statement

The authors have declared no competing interest.

