## Supplemental Table 1 for "Environmental RNAi-based reverse genetics in the predatory mite *Neoseiulus californicus*: towards improved methods of biological control"

**Table S1** Species list and GenBank accession numbers of species used in the evolutionary analysis of *N. californicus V-ATPase*.

| Species | Family | Order (suborder) | GenBank ID |
|---|---|---|---|
| *Sarcoptes scabiei* | Sarcoptidae | Acari (Astigmatina) | KPM04842.1 |
| *Dermatophagoides pteronyssinus* | Pyroglyphidae | Acari (Astigmatina) | XP_027205094.1 |
| *Ixodes scapularis* | Ixodidae | Acari (Ixodida) | XP_029849202.1 |
| *Galendromus occidentalis* | Phytoseiidae | Acari (Mesostigmata) | XP_003741079.1 |
| *Neoseiulus californicus* | Phytoseiidae | Acari (Mesostigmata) | QCS27658.1 |
| *Tropilaelaps mercedesae* | Laelapidae | Acari (Mesostigmata) | OQR76956.1 |
| *Varroa destructor* | Varroidae | Acari (Mesostigmata) | XP_022670784.1 |
| *Dinothrombium tinctorium* | Trombidiidae | Acari (Prostigmata) | RWS01677.1 |
| *Leptotrombidium deliense* | Trombiculidae | Acari (Prostigmata) | RWS24399.1 |
| *Tetranychus urticae* | Tetranychidae | Acari (Prostigmata) | XP_015785702.1 |
| *Aethina tumida* | Nitidulidae | Coleoptera | XP_019879320.1 |
| *Dendroctonus ponderosae* | Scolytinae | Coleoptera | XP_019764847.1 |
| *Harmonia axyridis* | Coccinellidae | Coleoptera | QCU54856.1 |
| *Hypothenemus hampei* | Curculionidae | Coleoptera | QEE14189.1 |
| *Leptinotarsa decemlineata* | Chrysomelidae | Coleoptera | XP_023012283.1 |
| *Tribolium castaneum* | Tenebrionidae | Coleoptera | XP_976188.1 |
| *Drosophila guanche* | Drosophilidae | Diptera | SPP82085.1 |
| *Bemisia tabaci* | Aleyrodidae | Hemiptera | XP_018897790.1 |
| *Apis mellifera* | Apidae | Hymenoptera | XP_016769524.1 |
| *Athalia rosae* | Tenthredinidae | Hymenoptera | XP_012267080.1 |
| *Cephus cinctus* | Cephidae | Hymenoptera | XP_015587567.1 |
| *Copidosoma floridanum* | Encyrtidae | Hymenoptera | XP_014208451.1 |
| *Dufourea novaeangliae* | Halictidae | Hymenoptera | XP_015436151.1 |
| *Eufriesea mexicana* | Apidae | Hymenoptera | OAD52578.1 |
| *Megachile rotundata* | Megachildae | Hymenoptera | XP_003700039.1 |
| *Microplitis demolitor* | Braconidae | Hymenoptera | XP_014297160.1 |
| *Nasonia vitripennis* | Pteromalidae | Hymenoptera | XP_001604685.1 |
| *Neodiprion lecontei* | Diprionidae | Hymenoptera | XP_015512625.1 |
| *Nylanderia fulva* | Formicidae | Hymenoptera | XP_029159858.1 |
| *Polistes dominula* | Vespidae | Hymenoptera | XP_015176675.1 |
| *Trichogramma pretiosum* | Trichogrammatidae | Hymenoptera | XP_014221633.1 |
| *Cryptotermes secundus* | Kalotermitidae | Isoptera | XP_023702715.1 |
| *Nasutitermes takasagoensis* | Termitidae | Isoptera | BAR72384.1 |
| *Reticulitermes flavipes* | Rhinotermitidae | Isoptera | AGO46410.1 |
| *Zootermopsis nevadensis* | Termopsidae | Isoptera | XP_021913539.1 |
| *Chilo suppressalis* | Crambidae | Lepidoptera | AXF48683.1 |
| *Galleria mellonella* | Pyralidae | Lepidoptera | XP_026752857.1 |
| *Spodoptera litura* | Noctuidae | Lepidoptera | XP_022826560.1 |
| *Frankliniella occidentalis* | Thripidae | Thysanoptera | XP_026274744.1 |
